# The DEPDC1 protein LET-99 is required for cortical stability and antagonizes branched actin formation to promote robust cytokinesis

**DOI:** 10.1101/2022.09.07.506988

**Authors:** Kari L. Price, Helen Lamb, Jocelyn V. Alvarado, Lesilee S. Rose

## Abstract

During cytokinesis, signals from the central spindle stimulate the accumulation of active RhoA-GTPase and thus contractile ring components at the equator, while the astral microtubules inhibit such components at the polar cortex. The DEPDC1 family protein LET-99 is required for furrow ingression in the absence of the central spindle signal, and for timely onset of furrowing even in the presence of the central spindle signal. Here we show that LET-99 works downstream or independently of RhoA-GTP and antagonizes branched F-actin and the Rac protein CED-10 to promote furrow initiation. This interaction with CED-10 is separable from LET-99’s function in spindle positioning. We also characterize a new role for LET-99 in regulating cortical stability, where LET-99 acts in parallel with the actomyosin scaffolding protein anillin, but LET-99 does not antagonize CED-10 in this case. We propose that LET-99 acts in a pathway that inhibits the Rac CED-10 to promote the proper balance of branched versus linear F-actin for cytokinesis, and that LET-99 also regulates another factor that contributes to cortical stability.

## Introduction

During cell division, after segregation of the duplicated chromosomes by the mitotic spindle, the mother cell is physically separated into daughter cells by the process of cytokinesis. In animal cells, cytokinesis is achieved via an actomyosin ring that pulls the cell membrane into a progressively constricting furrow. Proper timing and spatial positioning of the furrow are essential for the fidelity of chromosome segregation and the accurate partitioning of cytoplasmic components into the daughter cells (reviewed in (Glotzer, 2017; Green et al., 2012; Pollard and O’Shaughnessy, 2019; Verma et al., 2019).

The actomyosin contractile ring assembles in an equatorial band at the cortex in response to the localized recruitment and activation of the GTPase RhoA. When RhoA is activated by binding GTP (RhoA-GTP) it directly activates the diaphanous family of formins that promote the formation of linear actin filaments (linear F-actin). RhoA-GTP also activates Rho Kinase, which stimulates nonmuscle myosin II motor activity. In the contractile ring, myosin helps align bundles of linear F-actin, and the complex of contractile myosin and actin provides the driving force for constriction (Chircop, 2014; Glotzer, 2017; Green et al., 2012; Pollard and O’Shaughnessy, 2019; Verma et al., 2019).

The localization of RhoA-GTP and its effectors to the cell equator occurs in response to signals from the anaphase mitotic spindle. The overlapping, non-kinetochore-based microtubules of the central spindle recruit the centralspindlin complex, a heterotetramer of the kinesin protein MKLP1 and the Rac GTPase MgcRacGAP. The centralspindlin complex then promotes the accumulation of RhoA-GTP. In contrast, the long, centrosomal-emanating astral microtubules that touch the polar cortex inhibit the localization of RhoA-GTP from the poles (Glotzer, 2017; Green et al., 2012; Lewellyn et al., 2010; Mangal et al., 2018; Pollard and O’Shaughnessy, 2019; Verma et al., 2019; von Dassow, 2009; Werner and Glotzer, 2008; Werner et al., 2007).

In several systems, the additive effects of the astral and central spindle microtubules in stimulating cytokinesis can be observed at the level of the timing of cytokinesis or the extent of furrow ingression (Glotzer, 2017; Green et al., 2012; Lewellyn et al., 2010; Mangal et al., 2018; Pollard and O’Shaughnessy, 2019; Verma et al., 2019; von Dassow, 2009; Werner et al., 2007). For example, *C. elegans* embryos with changes in the proximity of the astral microtubules to the cortex exhibit delays in the timing of furrow initiation but cytokinesis completes. In embryos or cells missing a central spindle or depleted of the centralspindlin protein MKLP1, substantial ingression occurs but cytokinesis ultimately fails. Simultaneous perturbation of both centralspindlin activity and the astral signal results in a dramatic reduction or complete loss of furrow formation and ingression (Bringmann and Hyman, 2005; Dechant and Glotzer, 2003; Lewellyn et al., 2010; Mangal et al., 2018; Piekny and Glotzer, 2008; Tse et al., 2011; Werner and Glotzer, 2008; Werner et al., 2007; Zanin et al., 2013). Further, in *C. elegans* one-cell embryos, laser ablation of the spindle results in spatially separated astral versus central spindle induced furrows (Bringmann and Hyman, 2005) and mutants with a small misplaced spindle exhibit a centralspindlin-dependent furrow adjacent to the spindle, and an astral-mediated furrow away from the asters that does not require centralspindlin activity (Werner et al., 2007).

The depletion of several proteins significantly reduces furrowing only in conjunction with loss of centralspindlin components. Anillin (ANI-1 in *C. elegans*) is a scaffolding protein that binds myosin, actin, RhoA-GTP and other components, and is required for proper myosin organization during cytokinesis. Anillin binds active RhoA and is a downstream effector. Nonetheless, in some systems including *C. elegans*, the defects in cytokinesis observed upon loss of anillin are much more severe when the cell is dependent on astral inhibition to induce furrowing (Maddox et al., 2005; Piekny and Glotzer, 2008; Tse et al., 2011; van Oostende Triplet et al., 2014; Werner and Glotzer, 2008). This could reflect a greater need for anillin in large cells where the central spindle is far from cortex and/or a greater need for anillin when flow of actomyosin and RhoA-GTP away from the poles concentrates contractile ring components at the equator in response to astral inhibition (Reymann et al., 2016; von Dassow, 2009; Werner and Glotzer, 2008). On the other hand, the *C. elegans* protein NOP-1 activates RhoA, primarily as part of the aster-directed furrowing pathway, in parallel to centralspindlin (ZEN-4/CYK-4 in *C. elegans*) activation of RhoA at the equator. While *nop-1* mutants exhibit normal cytokinesis, *zen-4; nop-1* double mutant embryos do not exhibit any furrow ingression (Price and Rose, 2017; Tse et al., 2012).

Several regulators of spindle positioning during the first asymmetric division also enhance the furrow ingression defects of embryos without a central spindle or of *zen-4* mutants (Bringmann et al., 2007; Verbrugghe and White, 2007). Some of these regulators appear to affect actomyosin organization and furrowing indirectly, due to reduced spindle elongation or failure to displace the spindle posteriorly (Price and Rose, 2017; Werner et al., 2007). However, the role of the LET-99 protein in cytokinesis is genetically separable from its role in asymmetric division. For example, the PAR polarity proteins restrict the localization of LET-99 to a lateral posterior band in asymmetrically dividing cells (Tsou et al., 2002; Wu and Rose, 2007). However, the localization of LET-99 to the prospective cleavage furrow is determined by the spindle and occurs in both asymmetrically and symmetrically dividing cells, and loss of LET-99 enhances the furrow ingression defects of *zen-4* mutants in both types of cells as well (Bringmann et al., 2007; Price and Rose, 2017). Changes in the dynamics of cortical myosin have been reported for *let-99* mutants, and *let-99; zen-4* double mutants exhibit greatly reduced myosin accumulation at the furrow. In contrast, loss of *let-99* does not enhance the cytokinesis phenotype of *nop-1* mutants (Bringmann et al., 2007; Goulding et al., 2007; Price and Rose, 2017; Werner et al., 2007). However, it is not known if LET-99 affects furrowing by acting upstream of RhoA activation like NOP-1, if LET-99 functions with downstream RhoA-GTP effectors, or if rather LET-99 influences the actomyosin cytoskeleton via a separate pathway. Interestingly, the DEPDC1A and DEPDC1B family of proteins to which LET-99 is most related have a region with homology to Rho-GTPase activating proteins, and DEPDC1B has been shown to physically and genetically interact with Rac (Sendoel et al., 2014; Wu et al., 2015; Yang et al., 2014). These findings raise the possibility that LET-99 could genetically interact with Rac or other small G proteins during cytokinesis in *C. elegans*.

The small GTPases Cdc42 and Rac promote the formation of branched F-actin networks via the Arp2/3 nucleating complex. While depletion of Cdc42 and Rac typically do not cause failure of cytokinesis, studies in *C. elegans* and other systems has revealed that a proper balance of branched versus linear F-actin is critical for proper timing and mechanics of membrane ingression (Bastos et al., 2012; Canman et al., 2008; Chan et al., 2019; Chircop, 2014; Loria et al., 2012; Pal et al., 2020; Severson et al., 2002; Zhuravlev et al., 2017). In *C. elegans*, the Rac CED-10 negatively regulates cytokinetic furrowing: Hypomorphic mutations in *ced-10* can rescue the furrowing ingression defects of centralspindlin mutants. Similar results have been observed for depletion of Arp2 (ARX-2). Recent studies also found that strong depletion of ARX-2 affects timely furrow initiation and ingression rate; this effect can be rescued by partial depletion of the formin CYK-1, which promotes linear F-actin (Canman et al., 2008; Chan et al., 2019; Loria et al., 2012; Pal et al., 2020; Zhuravlev et al., 2017). The regulation of levels of branched versus linear F-actin are also critical for normal cell shape during cytokinesis and cell migration in many cell types, and ARX-2 and CED-10 have been implicated in regulating cortical stability in the early *C. elegans* embryo (Bustelo et al., 2007; Chircop, 2014; Loria et al., 2012; Roh-Johnson and Goldstein, 2009; Severson et al., 2002).

In this study, we further investigate LET-99’s role in regulating the actomyosin cytoskeleton. Our results indicate that LET-99 does not act upstream of activation of RhoA to promote cytokinetic furrowing, which together with prior work indicates that LET-99 is not specific to the astral inhibition pathway for cytokinesis. Rather, we show that LET-99 promotes timely furrow formation in a mechanism that acts antagonistically to branched F-actin and the Rac CED-10.

Further, we characterize a new role for LET-99 in promoting cortical stability. The cortical stability and cytokinesis defects of *let-99* embryos are enhanced by simultaneous loss of anillin. Surprisingly the cortical stability defects of *let-99* mutants are also enhanced, rather than suppressed by loss of CED-10. These results indicate that LET-99 is not in a simple linear pathway where it inhibits CED-10 to regulate both cytokinesis and cortical stability. From these and other data, we propose that LET-99 regulates the actomyosin cytoskeleton globally via two mechanisms: LET-99 inhibits branched F-actin formation and also influences other actomyosin regulators.

## Results

### LET-99 antagonizes branched F-actin filament formation to promote timely cytokinesis onset

The loss of LET-99 function enhances the cytokinetic furrow ingression defects of mutants missing the spindle midzone or the ZEN-4 component of the centralspindlin complex. In addition, embryos from *let-99* single mutant mothers (hereafter referred to as *let-99* embryos or mutants) were reported to have a delay in the onset of cytokinesis, even though cytokinesis completed (Bringmann et al., 2007; Price and Rose, 2017). We first confirmed that depletion of LET-99 by RNA interference (RNAi) results in a cytokinesis delay, so that this background could be used in further analyses of contractile ring components. The time from nuclear envelope breakdown (NEB) to furrowing onset (FO, defined by the first indentation of the membrane) was measured from DIC movies (Fig.1A, A’). We found that *let-99(RNAi)* embryos exhibited a delay in cytokinesis that was similar to that produced by the *let-99(dd17)* null mutant. Because loss of LET-99 also affects spindle positioning and elongation, which can indirectly affect furrowing, embryos with spindles misoriented at late anaphase were excluded from this and all subsequent analyses of furrowing kinetics (see Methods).

**Figure 1.**
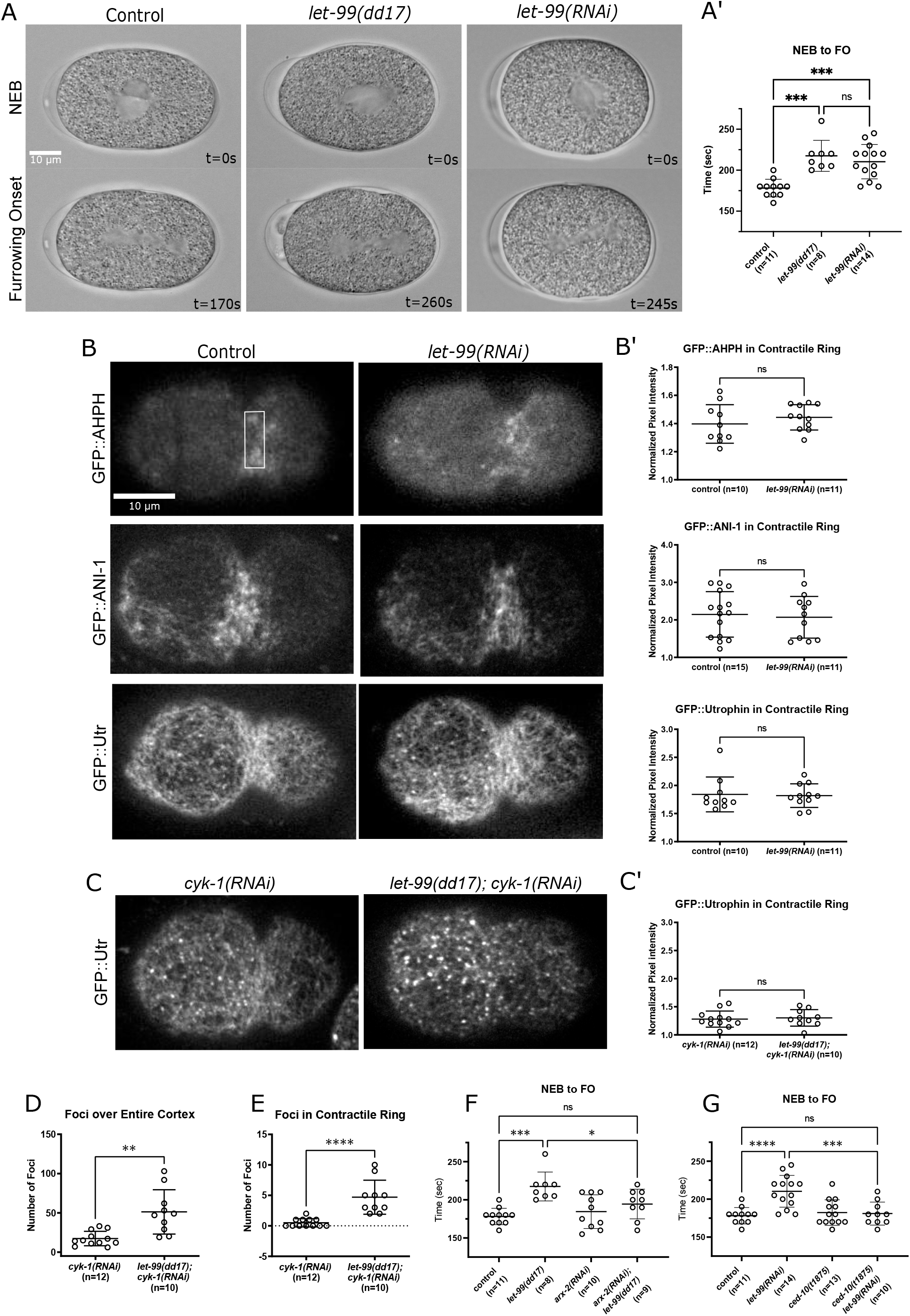
LET-99 antagonizes branched actin. (A) Images from DIC time-lapse microscopy illustrating nuclear envelope breakdown (NEB) and furrowing onset (FO) for the listed genotypes during the first cell division. Scale bar is 10 microns. Time is shown relative to NEB. Anterior is to the left in this and all figures. (A’) Quantification of time from NEB to FO. (B, C) Cortical images of representative embryos expressing components of the contractile ring as labeled. All images are scaled to the same pixel intensity values. Box in top left panel show the ROI of 20 × 60 pixels used to determine the average pixel intensity at the contractile ring for all quantifications shown in B’ and C’, where the normalized pixel intensity = value box on contractile ring/value of box on posterior cortex. (D, E) The number of foci were quantified either for the entire cortex or in an ROI of 20 × 60 pixels placed over the contractile ring. (F, G) Quantification of time from NEB to FO in the listed genotypes. Control was the wild-type N2 strain; control and dd17 data in A’ and F are the same. In all, error bars represent ± SD. Statistical significance was determined using Anova with multiple comparisons via the Sidak method for graphs in panels A’ and D-G. Unpaired student ttests were used for B’ and C’. ns = not significant (p > 0.05). See Supplementary Table S1 for specific P values.

To determine if LET-99 acts upstream of recruitment of activated RhoA, we examined embryos expressing GFP::AHPH, which contains the C-term half of anillin including its RhoA-GTP binding and plekstrin homology domains; this transgene has been validated as a sensor for active RhoA (Tse et al., 2012). Images were taken from a cortical view during the first cell division and enrichment in the contractile ring region was quantified (Fig. 1B, B’). No difference in the average fluorescence intensity between controls and *let-99(RNAi)* embryos was observed. The overall appearance of anillin labeled by GFP::ANI-1 (Tse et al., 2011), and the levels in the contractile ring were also comparable in control and *let-99(RNAi)* embryos (Fig. 1B, B’). These results suggest that LET-99 is not required for activation of RhoA and recruitment of anillin to the contractile ring.

RhoA-GTP promotes the accumulation of myosin and linear F-actin at the furrow. We previously reported lower levels of myosin at the furrow in single *let-99* mutants (Price and Rose, 2017). To examine F-actin localization during cytokinesis, we examined embryos expressing the GFP::Utrophin actin binding probe (GFP-Utr contains the calponin homology domain of Drosophila utrophin, which binds act (Tse et al., 2012). Total levels of F-actin at the furrow as assayed by GFP-Utr intensity were similar between control and *let-99(RNAi)* embryos (Fig. 1B, B’). However, we observed a qualitative increase in the number of small cortical foci of GFP:: Utr in *let-99* embryos. Cortical actin foci are indicative of branched actin filaments, because they are abolished by loss of ARX-2. In contrast depletion of CYK-1 lowers the levels of linear F-actin, making it easier to visualize the ARX-2 dependent foci (Chan et al., 2019; Shivas and Skop, 2012; Xiong et al., 2011). Thus to more easily visualize foci and quantify the effect of loss of LET-99 on branched F-actin, we depleted CYK-1 by RNAi in the GFP:: Utr-CH background, with or without *let-99(dd17*) and counted the number of foci. Both *cyk-1(RNAi)* and *cyk-1(RNAi); let-99(dd17)* embryos showed reduced levels of total GFP-Utr at the furrow compared to controls, as expected for knockdown of CYK-1 (Fig. 1C, C’). However, compared to *cyk-1(RNAi)* embryos, *cyk-1(RNAi); let-99(dd17)* embryos exhibited an increase in the number of GFP:: Utr-CH foci at the cleavage furrow as well as across the entire embryo cortex (Fig. 1D, E), These results indicate that there are increased levels of branched F-actin in *let-99* mutants.

The proper balance of Arp-2/3 dependent branched F-actin versus formin dependent linear F-actin is necessary for proper actomyosin organization during contractile ring assembly and thus for proper timing of furrowing onset, as indicated by the first shallow deformation of the membrane (Chan et al., 2019). We therefore tested whether reducing branched F-actin filaments, via RNAi of ARX-2, could rescue the furrowing delay of *let-99(dd17)* mutant embryos. The time to furrowing onset in *arx-2(RNAi)* embryos was similar to controls, consistent with prior results for partial depletion of ARX-2 (Chan et al., 2019). However, *arx-2(RNAi); let-99(dd17)* embryos exhibited a time to furrowing onset that was reduced compared to *let-99(dd17)* embryos (Fig. 1F). These results are consistent with a role for *let-99* in antagonizing branched F-actin filament formation during cytokinesis.

### LET-99 and CED-10/Rac have opposing roles during cytokinetic furrowing

In *C. elegans* embryos as in many systems, branched F-actin is promoted by both Rac and Cdc42 (CED-10 and CDC-42 in *C. elegans*)(Bustelo et al., 2007; Chircop, 2014). While neither *cdc-42* nor *ced-10* mutant embryos have overt cytokinesis phenotypes, several studies have shown that CED-10 antagonizes aspects of cytokinesis. For example, partial loss of function mutations in *ced-10* increases the extent of furrow ingression in embryos lacking centralspindlin function, as also seen for depletion of Arp2/3 (Canman et al., 2008; Jantsch-Plunger et al., 2000; Loria et al., 2012; Zhuravlev et al., 2017). The latter result is opposite to the effect of *let-99* mutations on furrow ingression in *centralspindlin* mutants (Price and Rose, 2017). We therefore tested whether loss of CED-10 function could rescue the delay in onset of furrowing exhibited by *let-99(RNAi)* embryos, using a maternal effect lethal allele that is a predicted null, *ced-10(t1875)* (Kinchen et al., 2005). In embryos from *ced-10(t1875)* mothers (hereafter referred to as *ced-10* embryos or mutants), there were no overt cytokinesis defects and the time from NEB to furrowing onset was comparable to controls (Fig. 1G), as reported previously for hypomorphic alleles or depletion of maternal CED-10 by RNAi. The mean time to furrowing onset in *let-99(RNAi) ced-10(t1875)* embryos was reduced compared to *let-99(RNAi)* embryos and was not significantly different than wild-type controls (Fig. 1G). Thus, similar to depletion of ARX-2, the loss of CED-10 restored furrow timing in *let-99(RNAi)* embryos.

Having established an antagonistic role between CED-10 and LET-99 for furrowing onset, we examined other aspects of cytokinesis. A balance of branched vs linear F-actin is also required for the timely transition from furrowing onset to the formation of a double-membraned furrow canal, also known as back-to-back membrane configuration, and for the rate of furrow ingression (Chan et al., 2019). To measure these aspects more accurately we utilized strains expressing endogenously tagged non-muscle myosin, NMY-2::mKate2, imaged in cross-section (Rehain-Bell et al., 2017)(Fig. 2A, see Methods). The time from NEB to furrowing onset in NMY-2::mKate2 control embryos was similar to that of wild-type embryos, and *let-99(dd17)* embryos showed a delay in furrowing onset, indicating that this background did not alter the cytokinesis phenotypes being scored (Fig. 2B). We found that in addition, *let-99(dd17)* embryos exhibited a longer time for the transition to back-to-back membrane formation compared to controls (Fig. 2C). The rate of furrow ingression in *let-99(dd17)* embryos was not statistically different from controls (Fig. 2D), consistent with a previous analysis of constriction of the entire furrow from an end on view (Bringmann et al., 2007). *ced-10(t1875)* embryos did not show any significant differences from controls in any of these measurements. In the *ced-10(t1875) let-99(dd17)* double mutant embryos, the time to furrowing onset was comparable to controls, and the time for back-to-back membrane formation was also significantly reduced in *ced-10(t1875) let-99(dd17)* double mutants compared to *let-99(dd17)* embryos (Fig. 2B, C). Together these data confirm that loss of CED-10 rescues the *let-99* mutant furrowing onset delay, and also reveal an antagonistic role between CED-10 and LET-99 in the transition from furrowing onset to back-to-back membrane formation.

**Figure 2.**
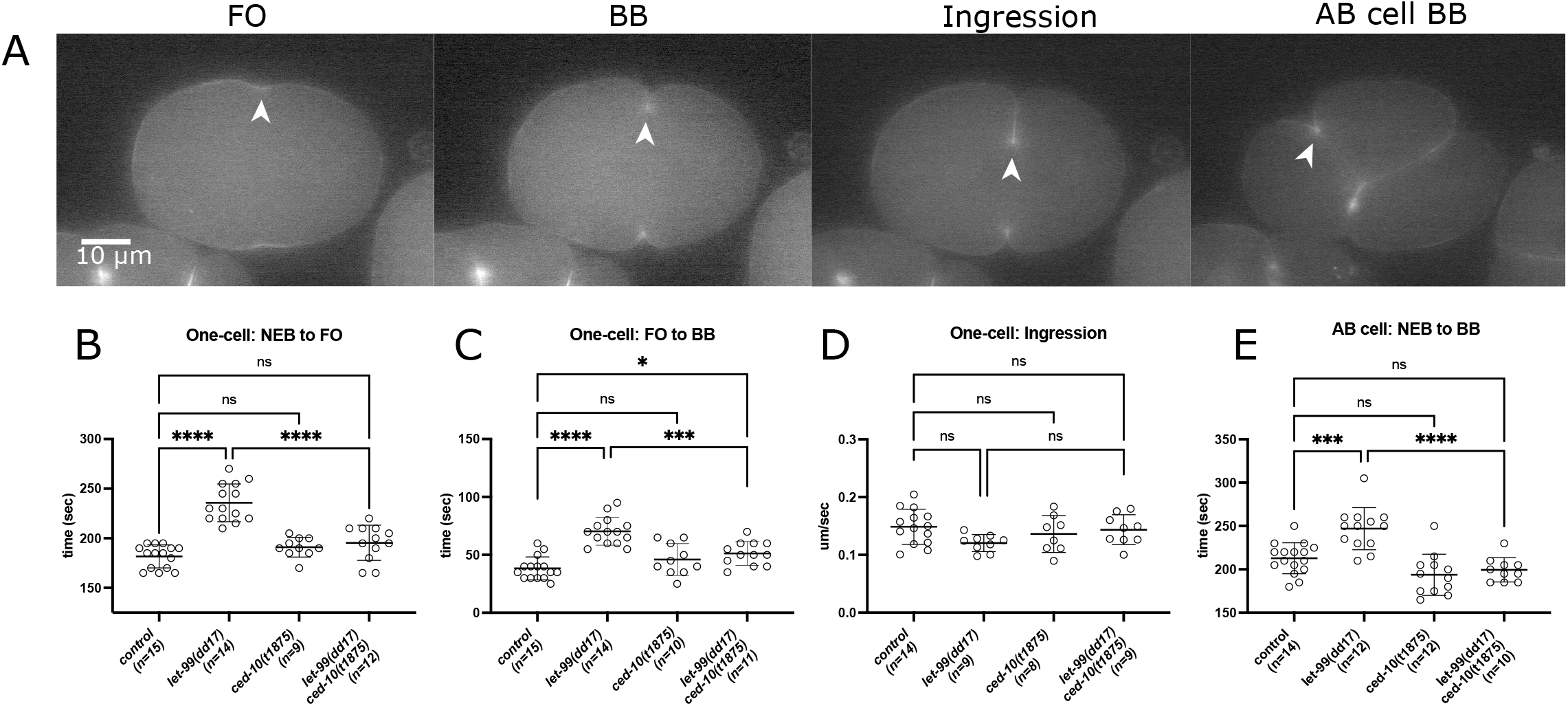
*let-99* embryos exhibit a delay in the transition from furrowing onset to back-to-back membrane formation that is rescued by loss of CED-10. (A) Images from fluorescent time-lapse microscopy of a representative NMY-2::GFP control embryo during the first two divisions. Scale bar is 10 microns. Time is shown relative to NEB and stages measured are shown: furrowing onset (FO), back-to-back membrane formation (BB) and furrow ingression (at BB plus 50 sec); white arrowhead identifies the leading furrow used for measurements (see Methods). (B-E) Quantification of times for stages as labeled in the one-cell embryo and in the AB cell of two-cell embryos. Error bars represent ± SD. Statistical significance was determined using Anova with multiple comparisons via the Sidak method. ns = not significant (p > 0.05). See Supplementary Table S2 for specific P values.

Loss of LET-99 also enhances the furrow ingression defects of centralspindlin deficient embryos in the AB cell; unlike the one-cell, the AB cell divides symmetrically and *let-99* mutants do not have defects in spindle elongation in this cell that could also impact cytokinesis (Price and Rose, 2017). We therefore examined furrow formation in the AB cell using the NMY-2::mKate background (Fig. 2A,E). The time from NEB to back-to-back membrane formation in the AB cell was slower in *let-99(dd17)* embryos compared to controls. In *ced-10(t1875) let-99(dd17)* double mutants, this timing was restored to control levels. These results confirm the antagonist relationship between LET-99 and CED-10 in cytokinetic furrowing and demonstrate that it is not specific to asymmetrically dividing cells.

### CED-10 does not affect spindle positioning in the one-cell embryo

Prior work showed that *let-99*’s enhancement of the cytokinesis defects of centralspindlin mutants did not correlate with defects in spindle position or elongation (Bringmann et al., 2007; Price and Rose, 2017). Interestingly however, CED-10 is required for spindle orientation in certain cells of 8-cell embryo and later in the vulval lineage (Cabello et al., 2010; Kishore and Sundaram, 2002). These observations raised the possibility that *let-99* and *ced-10* interact during spindle positioning. We therefore characterized spindle positioning in *ced-10(t1875)* and *ced-10(t1875) let-99(RNAi)* one-cell embryos in comparison to wildtype and *let-99(RNAi*) embryos.

In wild-type one-cell embryos, the male and female pronuclei meet in the posterior and then the nuclear-centrosome complex moves to the center of the embryo (centration); the complex also rotates so that the centrosomes are aligned on the polarized AP axis prior to NEB in most embryos (Figure 3A-D, Movie 1). The spindle is displaced posteriorly during metaphase and anaphase resulting in an unequal cleavage where the anterior AB cell is larger than the posterior P1 cell (Fig. 3A,B,E and Fig. 4A)(reviewed in (Rose and Gonczy, 2014)). In *let-99(RNAi)* or mutant embryos, there is a failure of centration, and variable rotation prior to NEB, which is accompanied by rocking of the nuclear centrosome complex. Nonetheless, the spindle aligns along the AP axis in most embryos prior to anaphase. Spindle elongation and displacement are both reduced but the overall spindle position is posterior and cleavage is unequal, because of the earlier centration defect (Fig. 3A-F, Movie 2, Fig. 4A) (Krueger et al., 2010; Rose and Kemphues, 1998; Tsou et al., 2002).

**Figure 3.**
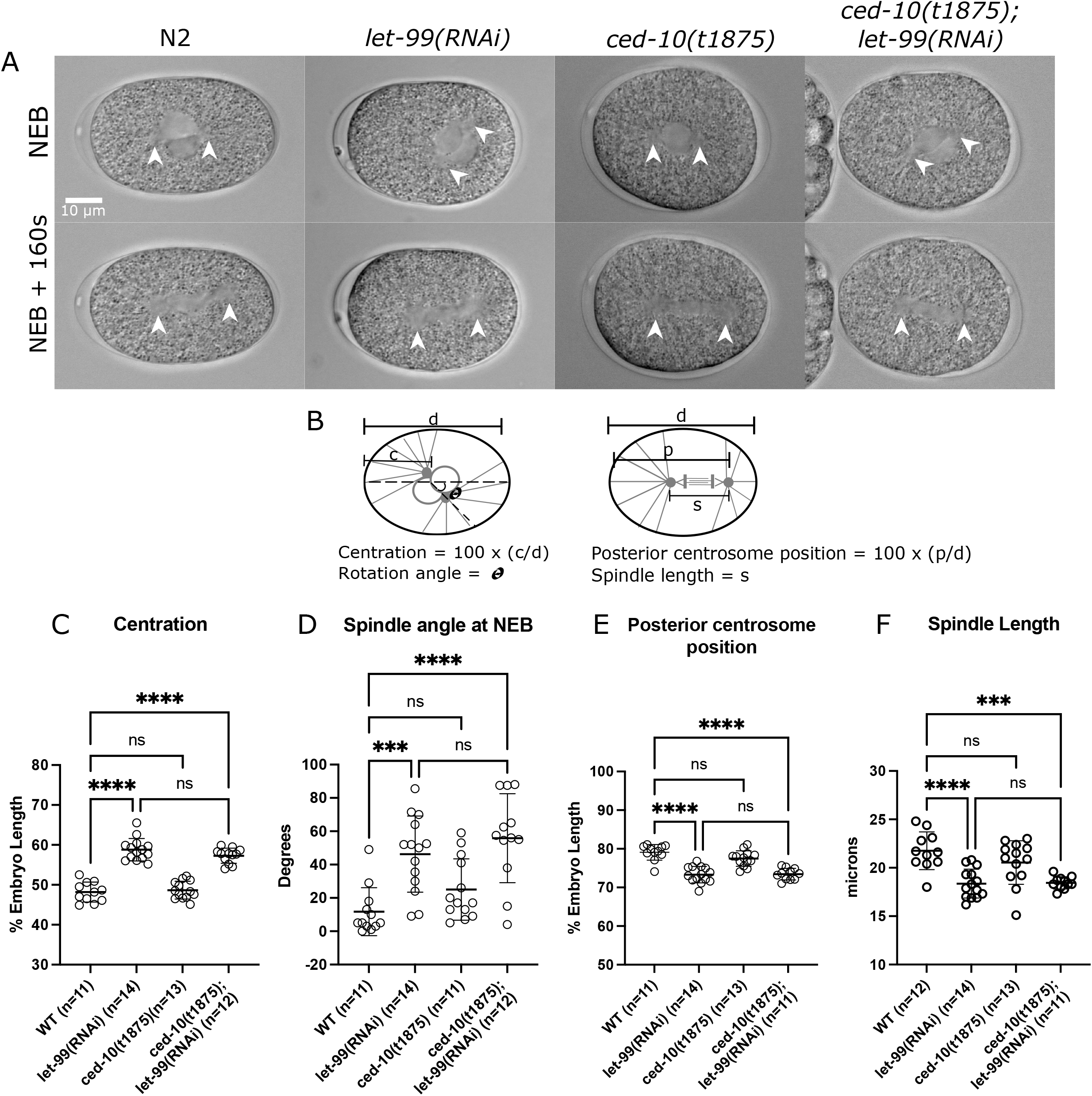
Loss of CED-10 does not affect one-cell spindle positioning. (A) Images from DIC microscopy of representative embryos with the listed genotypes at stages labeled. White arrowheads indicate the centrosomes. Anterior is to the left. Scale bar is 10 microns. (B) Schematic diagrams of measurements. Centration was determined by expressing the midpoint of the nuclear/centrosomal complex at NEB as a percentage of embryo length where anterior = 0% and posterior 100%. Rotation was measured as the angle between a line drawn along the anterior/posterior axis and a line connecting the centrosomes at NEB. As a readout for overall spindle placement, the position of the posterior centrosome as a percentage of % embryo length was measured at 160 sec after NEB. The length of the spindle was determined at 160 sec after NEB. Error bars represent ± SD. (C-F) Quantification of spindle positioning and elongation. Statistical significance was determined using Anova with multiple comparisons via the Sidak method. ns = not significant (p > 0.05). See Supplementary Table S3 for specific P values.

**Figure 4.**
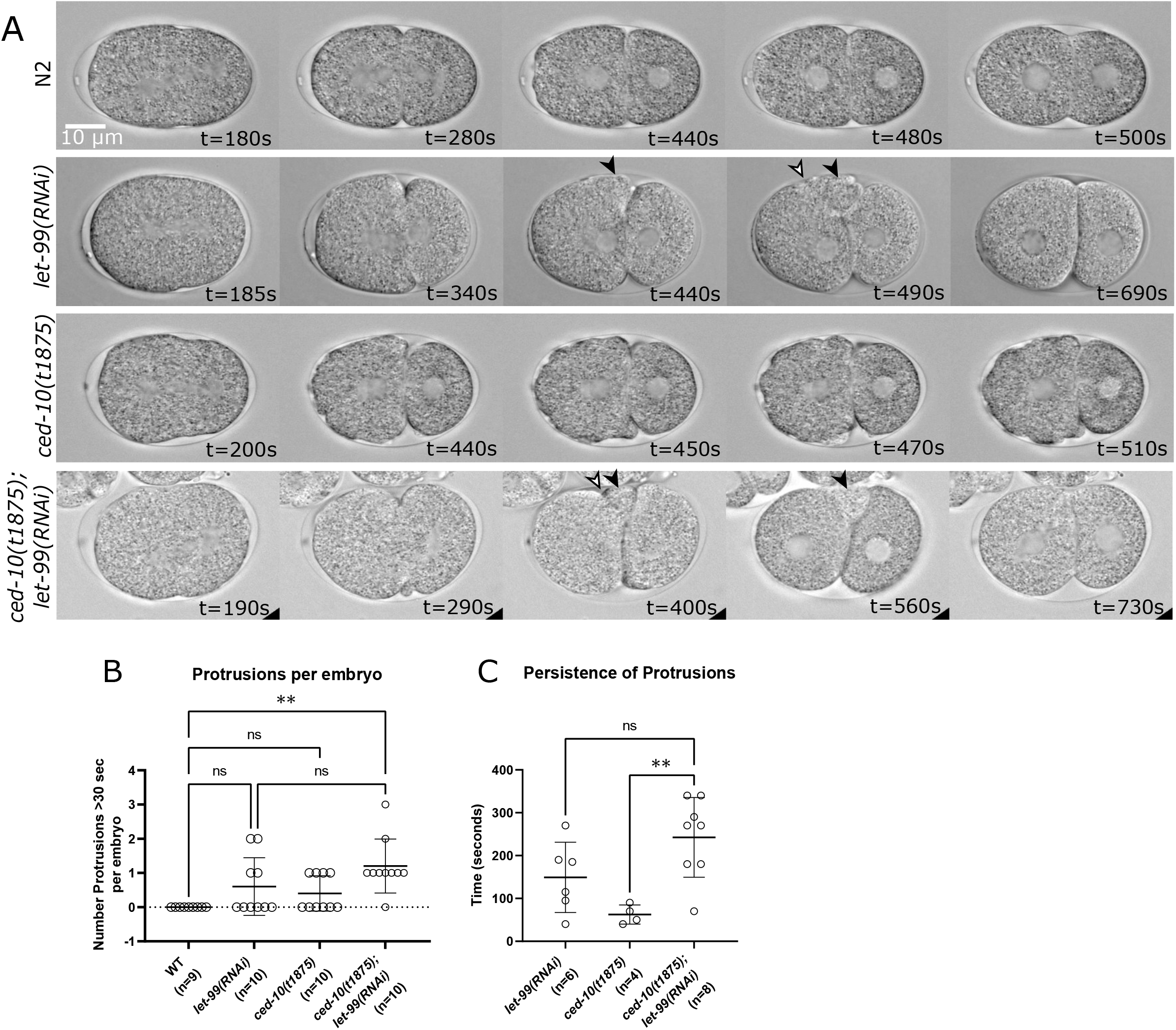
LET-99 and CED-10 enhance each other’s cortical instability phenotypes late in cytokinesis. (A) Stills from time-lapse DIC movies of the listed genotypes. Scale bar is 10 microns, anterior is the left. Black arrowheads mark persistent protrusions near the cytokinesis furrow; white arrowheads mark ectopic furrows that form in the protrusion. Time is shown relative to NEB = 0 seconds. Frames were chosen to illustrate the relative timing of different types of cortical activity described in the text, starting at furrowing onset or one frame after. (B) Quantification of the number of protrusions per embryo that lasted 40 sec or more. (C) Quantification of the persistence time of protrusions. Error bars represent ± SD. Statistical significance was determined using Anova with multiple comparisons via the Sidak method. ns = not significant (p > 0.05). See Supplementary Table S4 for specific P values.

In *ced-10(t1875)* embryos, the nuclear-centrosome complex centered and the spindle oriented onto the anterior-posterior axis before NEB, as observed in controls. The spindle then displaced back towards the posterior so that the one-cell divided unequally Fig. 3A-E, Movie 3, Fig. 4A). In addition, spindle elongation was comparable to controls. (Fig. 3F). The *ced-10(t1875) let-99(RNAi)* double mutant embryos exhibited nuclear centrosome rocking and defects in centration and nuclear rotation that were indistinguishable from those of *let-99(RNAi)* embryos (Fig.3A-D, Movie 4). The position of the posterior spindle pole was abnormal, and spindle length was reduced in the double mutant embryos, again comparable to *let-99(RNAi)* embryos (Fig. 3E,F). Thus, the loss of CED-10 does not suppress the spindle positioning or spindle elongation defects of *let-99* mutants. Overall, these results show that first, CED-10 has no apparent role in spindle positioning during the first division, and second, that the genetic interaction between *let-99* and *ced-10* mutations is specific to cytokinesis.

### LET-99 affects cortical stability

Although the normal timing of furrowing was restored in the *ced-10(t1875) let-99(RNAi)* embryos, many double mutants exhibited an ectopic furrow resulting in a wedge-like structure between the AB and P1 cells (Fig. 4A). Similar ectopic furrow structures were previously reported in *let-99* single mutants at the end of cytokinesis and were correlated with disorganization of the contractile ring as viewed end-on (Bringmann et al., 2007), but their genesis was not determined. We therefore examined the formation of these furrows in more detail. In *let-99(RNAi)* embryos, a single cytokinetic furrow formed and ingressed, similar to controls. However, 4/10 *let-99(RNAi)* embryos exhibited a protrusion of the AB cell membrane that extended toward the P1 cell and shifted the cell-cell contact region. We refer to this deformation as a half-wedge, because in all four of these embryos a second furrow emerged within the protrusion to produce the wedge structure (Fig. 4A, Movie 2). Although some protrusions in *let-99(RNAi)* embryos started during cytokinesis, the progression to the wedge occurred well after cytokinesis completion; the protrusions and extra furrows were also dynamic and resolved before the AB cell entered mitosis (Fig 4A, Movie 2). These observations indicate that *let-99* embryos do not initiate ectopic cytokinesis furrows at the start of cytokinesis, but rather suggest that *let-99* embryos have altered cortical activity of the AB cell.

Wild-type embryos have higher levels of cortical myosin at the anterior during cytokinesis of the one-cell and in the AB cell, and the AB cell cortex/membrane undulates during and after cytokinesis (Fujita and Onami, 2012; Loria et al., 2012; Munro et al., 2004; Ozugergin et al., 2022; Werner et al., 2007). Undulations of the AB cortex in control embryos occurred both in the polar regions and near the furrow, most lasting only 10-20 sec in one location (Movie 1). All *let-99(RNAi)* exhibited these typical rapid undulations of the AB cortex, but the furrow protrusions that developed into wedge structures lasted much longer, for more than 100 sec; two of the *let-99(RNAi)* embryos exhibited an additional protrusion of 40 sec or more (Fig. 4C). Thus, to compare the protrusive activity in single and double mutants, we quantified the number of “persistent” protrusions that lasted at least 40 sec in each embryo. Persistent protrusions near the furrow were also observed during or after cytokinesis in 4/10 *ced-10(t1875)* embryos, but these were of shorter duration than *let-99* protrusions, and in only one case did a wedge form (Fig. A-C). In general, there were more undulations of the entire AB cell membrane in *ced-10* embryos compared to controls (Fig. 4A Movie 3), consistent with prior observations of the anterior membrane during cytokinesis (Loria et al., 2012). Although the number of protrusions per embryo was similar in all mutant backgrounds, a higher proportion of *ced-10(t1875) let-99(RNAi)* double mutant embryos (9/10) exhibited persistent protrusions compared to either single mutant (Fig. 4B). In the subset of double mutant embryos which could be followed until late prophase of the AB cell, 4/5 embryos exhibited protrusions that progressed into wedges, and then ectopic furrows resolved (Fig. 4A, Movie 4). Some furrow protrusions in these embryos lasted longer compared to *let-99(RNAi)* embryos, but the overall mean persistence time was only significantly different from *ced-10* embryos (Fig. 4C). Similar results were observed in the NMY-2 background, where 92% of *ced-10(t1875) let-99(dd17)* (n= 12) exhibited a persistent wedge or half-wedge protrusion, compared to 41% of *let-99(dd17)* embryos (n = 17). Thus, this protrusion phenotype of *let-99* is enhanced rather than suppressed by simultaneous loss of *ced-10* and the data suggest that LET-99 regulates cortical stability.

We also observed a distinct type of cortical instability in *ced-10(t1875)* embryos, consisting of multiple very small bleb like structures (Movie 3). These structures were most apparent later in the two-cell stage compared to during cytokinesis, and could also be observed in the one-cell embryo at the end of the pseudocleavage stage through NEB. These small protrusions are similar to structures observed in *arx-2(RNAi)* embryos, which have been shown to be true blebs, that is, breaks in the cortical cytoskeleton through which membrane protrudes (Roh-Johnson and Goldstein, 2009; Severson et al., 2002). These small blebs were observed in *let-99(dd17) ced-10(t1875)* embryos as well. Together with the enhancement of the furrow protrusion phenotype, these results indicate that LET-99 and CED-10 are not in a simple linear pathway whereby LET-99 inhibits CED-10 to regulate both cytokinesis and cortical stability.

### *let-99; ani-1* double mutant embryos exhibit enhanced cortical instability

We previously showed *let-99(dd17);ani-1(RNAi);zen-4(RNAi)* embryos exhibit an even greater reduction in the extent of furrow ingression compared to *let-99(dd17);zen-4(RNAi)* or *ani-1(RNAi);zen-4(RNAi)* embryos, suggesting that LET-99 and ANI-1 act in parallel to promote cytokinetic furrowing (Price and Rose, 2017). We therefore tested whether loss of ANI-1 enhances the *let-99(dd17)* mutant phenotype even in the presence of ZEN-4, by examining the time to furrowing onset from DIC time-lapse movies. On average, *ani-1(RNAi)* embryos exhibited a normal time to furrowing onset. The *let-99(dd17);ani-1(RNAi)* embryos showed a delay which was significantly longer than in *let-99(dd17)* single mutants (Fig. 5A, F). However, *let-99(dd17); ani-1(RNAi)* embryos showed similar spindle positioning defects as the *let-99(dd17)* single mutants, and spindle elongation was not more defective in the double mutants (Fig. 5 B-E). These results show that LET-99 and ANI-1 act in parallel to promote timely furrowing onset, independently of effects on spindle elongation or position.

**Figure 5.**
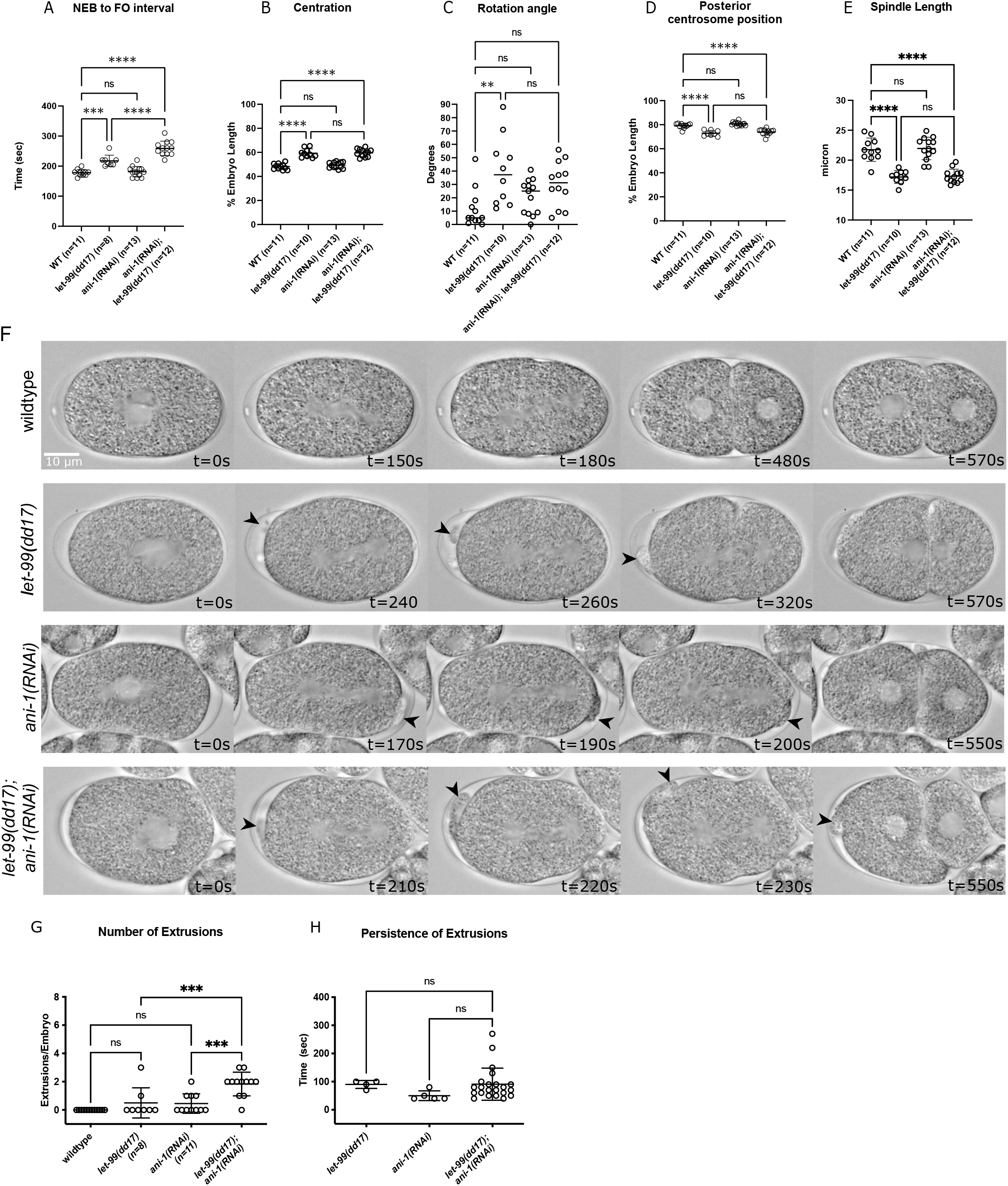
LET-99 and ANI-1 act in parallel to promote timely onset of furrow ingression and cortical stability. (A) Quantification of the time to furrowing onset. (B-E) Quantification of centration, rotation, posterior spindle pole position and spindle length measured as in Fig. 4. (F) Images from time-lapse DIC movies of the listed genotypes. Anterior is to the left and times (t) are relative to NEB = 0. Arrowheads mark extrusions. Scale bar, 10 *µ*m. (G) Quantification of the number of extrusions lasting 40 sec or more, per embryo. (H) Quantification of the persistence time of extrusions. Control and dd17 embryos are the same as in Fig. 1. Error bars represent ± SD. Statistical significance was determined using Anova with multiple comparisons via the Sidak method. ns = not significant (p > 0.05). See Supplementary Table S5 for specific P values.

The loss of anillin has been reported to result in protrusions at the furrow in Hela cells and cortical deformations have been observed in *C. elegans* one-cell embryos depleted of ANI-1 in combination with other cytokinesis regulators (Pacquelet et al., 2015; Price and Rose, 2017). We observed many protrusions near the furrow after cytokinesis in *ani-1(RNAi)* embryos and in *let-99(dd17);ani-1(RNAi)* embryos. In addition, we observed deformations of the cortex at anterior or posterior pole of the embryo during anaphase. We refer to these deformations as polar extrusions as they appear distinct from both furrow protrusions and the normal undulations of the AB membrane. A small number of *let-99(dd17)* single mutant embryos exhibited polar extrusions at the anterior (2/8, Fig. 5F,G; Movie 5; 0/10 *let-99(RNAi)* embryos exhibited extrusions). Several *ani-1(RNAi)* embryos (4/11) exhibited extrusions, all at the posterior pole (Fig. 5 F,G; Movie 6). The majority of *let-99(dd17); ani-1(RNAi)* double mutant embryos had polar extrusions and these could be at the anterior or pole posterior pole or both (11/12 embryos; Fig. 5 F,G; Movie 7). Strikingly, some extrusions in both the *ani-1(RNAi)* and *ani-1(RNAi); let-99(dd17)* embryos “traveled”, resolving at a distance from the site of initiation (Fig. 5F and Movie 7). Although the mean persistence time of the protrusions in the double mutant was not longer, the total number of extrusions per embryo was higher compared to the single mutants (Fig. 5G, H). Thus, simultaneous loss of LET-99 and ANI-1 results in enhanced instability of the cortex. Together with the analysis of *ced-10 let-99* double mutant embryos, these results identify a new role for LET-99 in regulating cortical stability. These data also provide evidence that LET-99 regulates the cortical cytoskeleton throughout the embryo, not just at the cytokinetic furrow.

## Discussion

The LET-99 protein was first identified as a regulator of spindle positioning during asymmetric division in *C. elegans*. Subsequent work revealed a redundant role for LET-99 in cytokinesis of both asymmetrically and symmetrically dividing cells. Although LET-99 accumulates asymmetrically at the cortex in response to polarity cues early in the cell cycle, the spindle promotes an equatorial enrichment of LET-99 at anaphase, independent of polarity (Bringmann et al., 2007; Price and Rose, 2017; Rose and Kemphues, 1998; Tsou et al., 2002). These and other data pointed to a role for LET-99 in regulating the actomyosin cytoskeleton during furrowing in a pathway separable from its role in spindle position. In this report, we further define the action of LET-99 during cytokinesis and identify a novel role for LET-99 in overall cortical stability.

Although *let-99* single mutants complete cytokinesis, they exhibit delays in furrowing onset, and here we identified a delay in the transition to the fully formed cleavage furrow. The loss of LET-99 also enhances the furrow ingression defects of embryos missing the central spindle or mutant for *zen-4*. In contrast, loss of LET-99 does not enhance the phenotype of *nop-1* mutants, and loss of LET-99 affects furrow ingression of the astral-mediated furrow in experiments where the astral and central spindle signals are spatially separated (Bringmann et al., 2007; Price and Rose, 2017; Werner et al., 2007). These genetic results initially placed LET-99 in the pathway for responding to astral inhibition signals. However, our recent work showed that the enrichment of LET-99 at the equator in anaphase can respond to signals from either the astral microtubules or the central spindle (Price and Rose, 2017). In the current study, we found that loss of LET-99 does not affect the contractile ring accumulation of a biosensor for active RhoA or full-length Anillin. These data together argue against models in which LET-99 is a specific component of the astral pathway that acts with NOP-1 to promote RhoA activation. Rather, LET-99 acts downstream from RhoA-GTP or in a separate pathway, and the genetic interactions reflect a greater need for LET-99 function when the actomyosin cytoskeleton is responding to astral signals, similar to the case for anillin. The effects of loss of anillin on myosin organization are distinct from those seen in LET-99 mutants (Adams et al., 1998; Bringmann et al., 2007; Price and Rose, 2017; Tse et al., 2011; Werner et al., 2007) and here we showed that *let-99; ani-1* double mutant embryos have enhanced defects in both cytokinesis and cortical stability. Thus, LET-99 and anillin regulate the actomyosin cytoskeleton through different mechanisms.

Our observations of increased branched F-actin foci in *let-99* embryos coupled with the rescue of *let-99’s* cytokinesis delays by reduction of Arp2 or the Rac CED-10 suggest that LET-99 has an antagonistic role to the formation of branched F-actin. The increased branched F-actin in the forming furrow of *let-99* mutant embryos may interfere directly with the alignment of linear F-actin and myosin in the contractile furrow (Canman et al., 2008; Descovich et al., 2018; Zhuravlev et al., 2017). LET-99 accumulates to highest levels at the equator. However, there are lower levels of LET-99 throughout the entire cortex (Price and Rose, 2017; Tsou et al., 2002), and the changes to actin foci observed were present throughout the cortex. Thus, the excess branched F-actin present in *let-99* mutants could also delay furrowing by increasing overall cortical stiffness outside the furrow or changing the dynamics of actomyosin globally (Taneja et al., 2020). In either case, loss of Arp2 or the Rac CED-10 in a *let-99* mutant would reduce branched F-actin, allowing for furrow formation with normal timing.

LET-99 could antagonize the formation of branched F-actin to promote furrowing through a number of different molecular mechanisms. There is cross-talk between Rho and Rac pathways in many processes, and the levels of Arp-dependent branched F-actin vs formin-dependent linear F-actin must be regulated precisely for proper cytokinesis. In several systems, depletion of Arp2/3 components or the small GTPase Rac increases the extent of furrow ingression in mutants with reduced RhoA-GTP activity caused by loss of centralspindlin. Similarly, although depletion of Arp2 doesn’t cause cytokinesis failure, it changes the timing of furrowing onset, back-to-back membrane formation, and furrow ingression. These effects can be ameliorated by simultaneous reduction of formin. Likewise defects caused by partial depletion of formin can be rescued by reduction of Arp2 (Bastos et al., 2012; Canman et al., 2008; Chan et al., 2019; Chircop, 2014; Glotzer, 2017; Lawson and Burridge, 2014; Loria et al., 2012; Pal et al., 2020; Pollard and O’Shaughnessy, 2019; Zhuravlev et al., 2017). A number of mechanisms, which are not mutually exclusive, have been implicated in this balancing act between branched and linear F-actin during cytokinesis in *C. elegans*. These include opposite effects of Rho and Rac on cortical rigidity, direct inhibition of Rac at the furrow by the centralsplindlin component MgcRacGAP (CYK-4), and global inhibition of the formin CYK-1 by Arp2/3 (Chan et al., 2019; Loria et al., 2012; Zhuravlev et al., 2017). LET-99 could be a new player in one of these pathways.

A mechanism in which LET-99 binds to Rac to directly inhibit its function is appealing because LET-99 is a member of the DEPDC1A and B family of proteins (Sendoel et al., 2014). This protein family contains an N-terminal DEP domain and a region with homology to Rho-GAP proteins near the C-terminus. Although the Rho-GAP motif is missing the catalytic residue for GAP activity in all family members, the domain could potentially bind small G proteins. Several studies have shown a genetic interaction between DEPDC1B and Rac or RhoA; one report showed that DEPDC1B binds Rac in vitro, and overexpression of DEPDC1B inhibited Rac activity and F-actin polymerization in Hela cells (Marchesi et al., 2014; Su et al., 2014; Wu et al., 2015; Yang et al., 2014). We have been unable to detect a consistent interaction between LET-99 and CED-10/Rac in vitro using recombinant proteins (unpublished results). This could reflect an in vivo requirement for other proteins to facilitate a LET-99 - Rac interaction. Alternatively, LET-99 could act indirectly via one of the pathways outlined above. Indeed, any model for LET-99 regulation of the actomyosin cytoskeleton must also take into account the observation that while loss of CED-10 suppressed the cytokinetic furrowing of *let-99* embryos, the double mutant embryos had enhanced cortical protrusive activity. Thus, we propose that LET-99 ultimately affects at least one other actomyosin cytoskeletal regulator or component in addition to branched F-actin. Precedence for this idea comes from known cross talk pathways for Rho and Rac. For example, Rho-kinase promotes myosin contractility but also activates negative regulators of Rac (Chircop, 2014; Lawson and Burridge, 2014).

LET-99 has been well-characterized as a key intermediate that acts downstream of PAR polarity cues to regulate spindle positioning in asymmetrically dividing cells. In that pathway, LET-99 acts to promote asymmetric pulling forces on the spindle, by inhibiting cortical localization of the Gα-dependent complex that recruits dynein (Park and Rose, 2008; Rose and Gonczy, 2014). Although the position of the astral microtubules is critical for myosin organization and flow in the astral mediated pathway for furrowing, prior work showed that LET-99’s requirement in furrowing does not correlate with spindle positioning or elongation defects (Bringmann et al., 2007; Price and Rose, 2017). Our current findings provide further evidence that LET-99 influences the actomyosin cytoskeleton in a mechanism independent of that used for spindle positioning. In particular, although CED-10 plays a role in spindle positioning in other cells (Cabello et al., 2010; Kishore and Sundaram, 2002), we found no role for CED-10 in the first division, and the *ced-10* null mutation neither enhanced or suppressed the spindle positioning defects of *let-99* mutant embryos. Similarly, the genetic enhancement between loss of LET-99 and ANI-1 is specific for cytokinesis and cortical instability. Thus, LET-99 appears to be a multifunctional protein that can act with either heterotrimeric or small G proteins in distinct mechanisms. Elucidating the molecular details of LET-99’s interactions will lead to a better understanding of the regulation of spindle positioning and the actomyosin cytoskeleton and should also give insight into the DEPDC1 family of proteins.

## Materials and Methods

### *C. elegans* strains

*C. elegans* strains were maintained on MYOB plates with *E. coli* OP50 as a food source (Brenner, 1974; Church et al., 1995). The following strains were used in this study.

| *Strain #* | *Genotype* |
| --- | --- |
| *N2* | *Wild type, Bristol strain* |
| *AZ244* | *unc-119(ed3) III; ruls57[pie-1::GFP::b-tubulin + unc-119(+)] V?* |
| *GE4047* | *ced-10(t1875)/ nT1[qIs50(pha::GFP) let- ?]* |
| *FM126* | *ltIs37[pie-1p::mCherry::his-58 + unc-119(+)] him-8(e1489) IV;*<br>*ruls57[pie-1::GFP::b-tubulin + unc-119(+)] V?* |
| *LP229* | *nmy-2(cp52[nmy-2::mKate2 + LoxP unc-119(+) LoxP]) I; unc-119(ed3) III* |
| *MG542* | *ltIs28 [pASM14; pie-1::GFP-TEV-STag::ani-1; unc-119 (+)];*<br>*ltIs37[pie-1p::mCherry::his-58; unc-119 (+)].* |
| *MG589* | *mgSi3[gfp::utrophin + Cbr-unc-119(+)]II; unc-119(ed3)III* |
| *MG617* | *mgSi5[gfp::ani-1(ahph) + Cbr-unc-119(+)].* |
| *RL175* | *unc-22(e66) let-99(dd17) / nT1[unc-? (n754d) let-?].* |
| *RL231* | *unc-22(e66) let-99(dd17) / nT1[qIs51(pha::GFP) let-?] IV* |
| *RL356* | *ced-10(t1875) dpy-20(e1282) unc-22(e66) let-99(dd17) /*<br>*nT1 [qIs51(pha::GFP)] V* |
| *RL263* | *ced-10(t1875)/nT1; qIs50[pha::GFP]; (ruls57 [GFP::tubulin; unc-119(+)].* |
| *RL359* | *ced-10(t1875) dpy-20(e1282) /nT1 [qIs51(pha::GFP)]* |
| *RL361* | *mgSi3[cb-unc-119(+) GFP::Utrophin] II;*<br>*ltIs37[pie-1p::mCherry::his-58 + unc-119(+)] him-8(e1489) IV* |
| *RL369* | *nmy-2(cp52[nmy-2::mkate2 + LoxP unc-119(+) LoxP]) I; unc-119(ed3) III;*<br>*ced-10(t1875) dpy-20(e1282) IV /nT1 [qIs51(pha::GFP)] V* |
| *RL408* | *nmy-2(cp52[nmy-2::mKate2]) I; let-99(dd17) unc-22(e66)/ nT1*<br>*[qIs51(pha::GFP)] IV* |
| *RL409* | *nmy-2(cp52[nmy-2::mKate2]) I; let-99(dd17) unc-22(e66) ced-10(t1875) dpy-*<br>*20/ nT1 [qIs51(pha::GFP)] V* |

### RNAi

RNAi was performed by feeding (Timmons et al., 2001). Constructs used to induce RNAi were obtained from the Ahringer RNAi library (Kamath et al., 2003)and include *ani-1 (III-6E18), let-99 (IV-6P07), cyk-1(III-8B02), arx-2(V-7M13)*. The extent of RNAi penetrance was determined from examining embryos for published strong loss of function phenotypes as follows: 1) *ani-1:* lack of pseudocleavage (Maddox et al., 2005), 2) *let-99:* severe nuclear-centrosome rocking, failure of centration in the one-cell, and failure of P1 nuclear rotation (Tsou et al., 2002), 3) *cyk-1:* weak pseudocleavage and reduced F-actin in the cleavage furrow (Ding et al., 2017; Swan et al., 1998) and 4) *arx-2:* increased number of anterior, cortical blebs (Severson et al., 2002). In cases where a double mutant background might suppress one mutant phenotype, the RNAi strength was also assessed by examining control embryos treated in parallel.

### Microscopy

Worms were dissected in egg buffer (25 mM HEPES, pH 7.4, 120 mM NaCl, 48 mM KCl, 2 mM CaCl_2_, MgCl_2_) and embryos were mounted on 2% agarose pads in the same buffer and imaged. For the analyses of furrowing onset, spindle positioning, cortical stability in Figures 1 and 3-5, embryos were examined in differential interference contrast (DIC) using an Olympus BX60 compound microscope, using a PlanApo N 60x, 1.42 numerical aperture oil-immersion objective. Single plane time-lapse images were acquired every 5 or 10 seconds using a Hamamatsu Orca 12-bit digital camera and Micromanager software (Edelstein et al., 2001). For analysis of cytokinesis in embryos expressing NMY-2::mKate-2 in Figure 2, embryos were mounted as above and imaged with the same system; images were taken every 5 sec in bright field through NEB, then in widefield fluorescence from metaphase through cytokinesis completion. Room temperature during imaging was 23-25°C.

To image GFP::AHPH, GFP::ANI-1 and GFP::Utr from a cortical view, embryos were mounted as above and imaged using the spinning disk module of a Marianas SDC Real-Time 3D Confocal-TIRF microscope (Intelligent Imaging Innovations, 3i) fit with a Yokogawa spinning disk head, a 63x 1.3 numerical aperture oil-immersion objective, and EMCCD camera. Acquisition was controlled by Slidebook 6 software (3i Incorporated). Single focal planes images were acquired first in midfocal plan to score NEB and anaphase onset timing; the focal plane was then moved to a cortical view during anaphase, and images were obtained every 3 sec at 100ms exposure. Room temperature during imaging was 24-25°C. Raw images were exposed and scaled with the same parameters for each experiment, and all images were processed using Fiji software (Schindelin et al., 2012).

### Analysis of cytokinetic furrowing

Cytokinetic furrowing rate measurements for both the P_0_ cell and the AB cell were taken relative to nuclear envelope breakdown (NEB). For all analyses, NEB was scored under DIC or brightfield conditions, NEB was scored as the first frame in which the pronuclei or nucleus began to shrink in size. The time of furrowing onset was scored as the frame in which the first cortical indentation (aka shallow deformation) that proceeded to furrow formation was observated. Membrane ingression in the one-cell C. *elegans* embryos is asymmetric: although furrowing onset occurs fairly uniformly around the circumference, one furrow (the leading furrow) transitions to the back-to-back membrane (BB) configuration faster and ingresses farther; the other furrow forms with more variable timing (Maddox et al., 2007). Thus measurements of back-to-back membrane and furrow ingression are only reported for the leading furrow and could only be scored accurately in fluorescence in the NMY-2::mKate2 background. BB formation was scored as the first frame where the membranes changed from a “V” configuration to back-to-back membranes where a furrow tip was visible. To measure ingression rate, the Manual Tracking plugin in Fiji software (Schindelin et al., 2012) was used to track the movement of the furrow tip until cytokinesis completion. The tracked distance during the first 50 seconds after BB was divided by time to give an average velocity. For this and all analyses, a sample of at least n = 10 embryos was obtained for each genotype. However, in some cases embryos were excluded so the final n for each genotype varies. In particular, because misoriented spindles at late anaphase could affect signaling from the spindle to the cortex and impact cytokinesis separately from LET-99’s function in furrowing, only embryos in which the spindle was aligned on the anterior-posterior axis by mid-anaphase were quantified for cytokinesis assays. In addition, we observed that a small number of *let-99(dd17)* embryos appeared to have a much slower cell cycle overall, where the time for all stages was approximately 1.5x that of other *let-99* embryos and controls, and the spindle at NEB plus 160s was clearly still in metaphase instead of anaphase. We thus used failure to progress into anaphase in a timely fashion as a criterion to exclude such embryos from all single and double mutant analyses using *let-99(dd17)*. This slow phenotype was not observed after RNAi of *let-99*. Furrowing measurements were assembled in excel and then imported into Prism for statistical analysis and graphing.

### Analysis of cortical contractile ring components

To measure enrichment of contractile ring components for Fig.1B, B’, mean pixel intensity values corresponding to a 20 pixel x 60 pixel region of interest placed over the forming furrow were obtained and normalized to the signal on the posterior side of the contractile ring. For measurements of GFP::ANI-1, measurements were made at ∼90 sec after anaphase onset, as determined by monitoring mCherry: H2B labeling of the DNA in cross section. The GFP::AHPH and GFP::Utrophin strains did not have a DNA marker; for these, measurements were made within 20-50 sec after furrowing onset, which is similar in timing to 90 sec after anaphase onset. To determine the number of GFP::Utrophin foci at the cortex in *cyk-1* backgrounds (Fig.1 C, C’), images were first thresholded in ImageJ using the Shanbhag image thresholding plugin with a minimum value of two pixels required to define a focus. Foci were then counted using the Particle Analysis tool in Fiji (Schindelin et al., 2012) either over the entire cortex or in a 20 × 60 pixel ROI over the furrow region. Data were compiled in Microsoft Excel and then exported into Prism for statistical analysis.

### Analysis of spindle positioning and elongation

Parameters of nuclear and spindle positioning were measured in Image J (https://imagej.nih.gov/ij/) using the line and angle tools, as shown in Fig. 3 and the corresponding text. Centration and rotation were measured at NEB. The position of the posterior spindle pole and spindle length were measured at NEB plus 160 sec, when anaphase spindles have elongated in controls embryos, but furrowing has not initiated. Measurements were compiled in Microsoft Excel and then exported into Prism for statistical analysis.

### Analysis of cortical stability

Protrusions were measured manually by scoring the first frame where a deformation of the cortex was observed (start) and the first frame where the protrusion had completely disappeared (resolution). The total time was calculated as (resolution frame – start frame) x frame rate. In controls, undulations of the entire cortex and protrusions near the furrow typically lasted only 10 or 20 sec, but an occasional deformation lasted 30 sec. Therefore, only protrusions lasting 40 sec or longer were scored as persistent. In all *let-99* embryos that were imaged through the second division, furrow protrusions began during one-cell cytokinesis or within 90 sec of furrowing completion, and all protrusions resolved before the AB cell underwent NEB. For scoring other genotypes, movies that lasted at least 400sec after NEB were used to score the the number of protrusions that initiated. For quantifying mean persistence time, only protrusions that resolved before the movie ended were used. The persistence of polar extrusions were measured in a similar manner. Data were assembled in excel and then imported into Prism for statistical analysis and graphing.

## Acknowledgements

We thank members of the Rose and McNally labs for helpful discussions. We are grateful to Michael Glotzer, Bob Goldstein, Amy Maddox and Frank McNally for provided strains. We also thank the *Caenorhabditis* Genetics Center, which is funded by the National Institutes of Health Office of Research Infrastructure Programs (P40 OD010440) for strains. The 3i Marianas spinning disk confocal used in this study was purchased using a National Institutes of Health Shared Instrumentation Grant [1S10RR024543-01]. We thank the MCB Light Microscopy Imaging Facility, which is a UC-Davis Campus Core Research Facility, for the use of this microscope. We are grateful to Kathie Urrutia-Paniagua for media preparation. This work was funded by the National Institutes of Health Grant [R01GM68744] and the National Institute of Food and Agriculture [CA-D*-MCB-6239-H]. Student support was also provided by the Floyd and Mary Schwall Dissertation Year Fellowship [KLP], the UC Davis BMCDB Graduate Group [KLP, HL] and a National Institutes of Health training grant [T32 GM 007377].

## Figure Legends

Tables S1-S5 and Movie legends are in the Supplemental Materials File.

