## Supplemental Tables and Movie legends for "The DEPDC1 protein LET-99 is required for cortical stability and antagonizes branched actin formation to promote robust cytokinesis"

Table S1. Statistics corresponding to data presented in Fig. 1

**Fig. 1A'  NEB – Furrowing Onset interval (seconds)**

| Condition | Mean | SD | n | ANOVA, Šídák * | |
|---|---|---|---|---|---|
| | | | | **comparison** | **p-value** |
| control | 178.2 | 10.8 | 11 | N/A | N/A |
| let-99(dd17) | 217.5 | 18.9 | 8 | vs WT | 0.0001 |
| let-99(RNAi) | 210.4 | 21.0 | 14 | vs WT | 0.0003 |
| | | | | vs let-99(dd17) | 0.6376 |

**Fig. 1B'  Contractile ring (normalized pixel intensity)**

| Condition | Mean | SD | n | Unpaired t test with Welch's correction | |
|---|---|---|---|---|---|
| | | | | **comparison** | **p-value** |
| GFP::AHPH control | 1.40 | 0.14 | 10 | N/A | N/A |
| GFP::AHPH let-99(RNAi) | 1.44 | 0.09 | 11 | vs control | 0.3708 |
| GFP::ANI-1 control | 2.15 | 0.61 | 15 | N/A | N/A |
| GFP::ANI-1 let-99(RNAi) | 2.07 | 0.56 | 11 | vs control | 0.7407 |
| GFP::Utr control | 1.84 | 0.31 | 10 | N/A | N/A |
| GFP::Utr let-99(RNAi) | 1.82 | 0.21 | 11 | vs control | 0.8551 |

**Fig. 1C'  GFP::Utrophin in contractile ring (normalized pixel intensity)**

| Condition | Mean | SD | n | Unpaired t test with Welch's correction | |
|---|---|---|---|---|---|
| | | | | **comparison** | **p-value** |
| cyk-1(RNAi) | 1.28 | 0.14 | 12 | N/A | N/A |
| let-99(dd17);cyk-1(RNAi) | 1.30 | 0.15 | 10 | vs cyk-1(RNAi) | 0.7403 |

**Fig. 1D  Foci over entire cortex (number)**

| Condition | Mean | SD | n | Unpaired t test with Welch's correction | |
|---|---|---|---|---|---|
| | | | | **comparison** | **p-value** |
| cyk-1(RNAi) | 17.3 | 9.2 | 12 | N/A | N/A |
| let-99(dd17);cyk-1(RNAi) | 51.2 | 28.2 | 10 | vs cyk-1(RNAi) | 0.0041 |

**Fig. 1E  Foci in contractile ring (number)**

| Condition | Mean | SD | n | Unpaired t test with Welch's correction | |
|---|---|---|---|---|---|
| | | | | **comparison** | **p-value** |
| cyk-1(RNAi) | 0.5 | 0.6 | 12 | N/A | N/A |
| let-99(dd17);cyk-1(RNAi) | 4.7 | 2.8 | 10 | vs cyk-1(RNAi) | <0.0001 |

**Fig. 1F, G  NEB – Furrowing Onset interval (seconds)**

| Condition | Mean | SD | n | ANOVA, Šídák * | |
|---|---|---|---|---|---|
| | | | | **comparison** | **p-value** |
| control | 178.2 | 10.8 | 11 | N/A | N/A |
| let-99(dd17) | 217.5 | 18.9 | 8 | vs WT | 0.0002 |
| arx-2(RNAi) | 184.5 | 22.0 | 10 | vs WT | 0.8931 |
| let-99(dd17);arx-2(RNAi) | 194.4 | 19.4 | 9 | vs WT | 0.1952 |
| | | | | vs let-99(dd17) | 0.0499 |
| let-99(RNAi) | 210.4 | 21.0 | 14 | vs WT | <0.0001 |
| ced-10(t1875) | 182.3 | 16.9 | 13 | vs WT | 0.9596 |
| ced-10(t1875);let-99(RNAi) | 181.0 | 15.2 | 10 | vs ced-10(t1875) | 0.9922 |
| | | | | | 0.0005 |

* Ordinary one-way ANOVA followed by Šídák's multiple comparisons test

Table S2. Statistics corresponding to data presented in Fig. 2

| **Fig. 2B NEB – Furrowing Onset interval (seconds)** | | | | | |
|---|---|---|---|---|---|
| **Condition** | **Mean** | **SD** | **n** | **Ordinary one-way ANOVA followed by Šídák's multiple comparisons test** | |
| | | | | **comparison** | **p-value** |
| control | 181.7 | 11.6 | 15 | N/A | N/A |
| let-99(dd17) | 235.7 | 19.0 | 14 | vs WT | <0.0001 |
| ced-10(t1875) | 191.0 | 9.9 | 10 | vs WT | 0.4547 |
| ced-10(t1875);let-99(dd17) | 195.4 | 17.7 | 12 | vs WT | 0.0938 |
| | | | | vs let-99(dd17) | <0.0001 |
| **Fig. 2C Furrowing – Back-to-Back interval (seconds)** | | | | | |
| **Condition** | **Mean** | **SD** | **n** | **Ordinary one-way ANOVA followed by Šídák's multiple comparisons test** | |
| | | | | **comparison** | **p-value** |
| control | 38.3 | 9.9 | 15 | N/A | N/A |
| let-99(dd17) | 70.4 | 12.0 | 14 | vs WT | <0.0001 |
| ced-10(t1875) | 46.0 | 13.7 | 10 | vs WT | 0.3613 |
| ced-10(t1875);let-99(dd17) | 51.3 | 10.3 | 12 | vs WT | 0.0209 |
| | | | | vs let-99(dd17) | 0.0004 |
| **Fig. 2D Ingression rate (microns/second)** | | | | | |
| **Condition** | **Mean** | **SD** | **n** | **Ordinary one-way ANOVA followed by Šídák's multiple comparisons test** | |
| | | | | **comparison** | **p-value** |
| control | 0.149 | 0.030 | 14 | N/A | N/A |
| let-99(dd17) | 0.121 | 0.015 | 9 | vs WT | 0.0753 |
| ced-10(t1875) | 0.136 | 0.032 | 8 | vs WT | 0.7608 |
| ced-10(t1875);let-99(dd17) | 0.144 | 0.026 | 9 | vs WT | 0.9863 |
| | | | | vs let-99(dd17) | 0.2777 |
| **Fig. 2E AB cell NEB – Back-to-Back interval (seconds) in mKate2::NMY-2 background** | | | | | |
| **Condition** | **Mean** | **SD** | **n** | **Ordinary one-way ANOVA followed by Šídák's multiple comparisons test** | |
| | | | | **comparison** | **p-value** |
| control | 212.8 | 17.9 | 14 | N/A | N/A |
| let-99(dd17) | 246.9 | 24.3 | 12 | vs WT | 0.0002 |
| ced-10(t1875) | 193.8 | 23.8 | 12 | vs WT | 0.0740 |
| ced-10(t1875);let-99(dd17) | 199.5 | 14.0 | 10 | vs WT | 0.3860 |
| | | | | vs let-99(dd17) | <0.0001 |

Table S3. Statistics corresponding to data presented in Fig. 3

| Fig. 3C Centration at NEB (% Embryo Length) | | | | | |
|---|---|---|---|---|---|
| Condition | Mean | SD | n | Ordinary one-way ANOVA followed by Šídák's multiple comparisons test | |
| | | | | comparison | p-value |
| WT | 48.2 | 2.5 | 11 | N/A | N/A |
| let-99(RNAi) | 58.8 | 2.8 | 14 | vs WT | <0.0001 |
| ced-10(t1875) | 48.7 | 2.2 | 13 | vs WT | 0.9805 |
| ced-10(t1875);let-99(RNAi) | 57.3 | 1.8 | 12 | vs WT | <0.0001 |
| | | | | vs let-99(RNAi) | 0.3869 |

| Fig. 3D Spindle angle at NEB (degrees) | | | | | |
|---|---|---|---|---|---|
| Condition | Mean | SD | n | Ordinary one-way ANOVA followed by Šídák's multiple comparisons test | |
| | | | | comparison | p-value |
| WT | 11.8 | 14.4 | 11 | N/A | N/A |
| let-99(RNAi) | 46.3 | 22.8 | 14 | vs WT | 0.0005 |
| ced-10(t1875) | 25.1 | 18.4 | 11 | vs WT | 0.4045 |
| ced-10(t1875);let-99(RNAi) | 55.9 | 26.7 | 12 | vs WT | <0.0001 |
| | | | | vs let-99(RNAi) | 0.6904 |

| Fig. 3E Posterior centrosome position at NEB + 160s (% Embryo Length) | | | | | |
|---|---|---|---|---|---|
| Condition | Mean | SD | n | Ordinary one-way ANOVA followed by Šídák's multiple comparisons test | |
| | | | | comparison | p-value |
| WT | 79.1 | 2.0 | 11 | N/A | N/A |
| let-99(RNAi) | 73.3 | 2.1 | 14 | vs WT | <0.0001 |
| ced-10(t1875) | 77.5 | 2.0 | 13 | vs WT | 0.1828 |
| ced-10(t1875);let-99(RNAi) | 73.5 | 1.5 | 11 | vs WT | <0.0001 |
| | | | | vs let-99(RNAi) | 0.9994 |

| Fig. 3F Spindle length at NEB + 160s (microns) | | | | | |
|---|---|---|---|---|---|
| Condition | Mean | SD | n | Ordinary one-way ANOVA followed by Šídák's multiple comparisons test | |
| | | | | comparison | p-value |
| WT | 21.8 | 1.9 | 11 | N/A | N/A |
| let-99(RNAi) | 18.4 | 1.5 | 14 | vs WT | <0.0001 |
| ced-10(t1875) | 20.6 | 2.3 | 13 | vs WT | 0.3200 |
| ced-10(t1875);let-99(RNAi) | 18.5 | 0.6 | 11 | vs WT | 0.0002 |
| | | | | vs let-99(RNAi) | >0.9999 |

Table S4. Statistics corresponding to data presented in Fig. 4

| **Fig. 4B Protrusions per embryo (number)** | | | | | |
|---|---|---|---|---|---|
| **Condition** | **Mean** | **SD** | **n** | **Ordinary one-way ANOVA followed by Šídák's multiple comparisons test** | |
| | | | | **comparison** | **p-value** |
| WT | 0 | 0 | 9 | N/A | N/A |
| let-99(RNAi) | 0.6 | 0.8 | 10 | vs WT | 0.1834 |
| ced-10(t1875) | 0.4 | 0.5 | 10 | vs WT | 0.5553 |
| ced-10(t1875);let-99(RNAi) | 1.2 | 0.8 | 10 | vs WT | 0.0010 |
| | | | | vs let-99(RNAi) | 0.1640 |
| **Fig. 4C Persistence of protrusions (seconds)** | | | | | |
| **Condition** | **Mean** | **SD** | **n** | **Ordinary one-way ANOVA followed by Šídák's multiple comparisons test** | |
| | | | | **comparison** | **p-value** |
| let-99(RNAi) | 149.2 | 81.9 | 6 | N/A | N/A |
| ced-10(t1875) | 62.5 | 22.2 | 4 | N/A | N/A |
| ced-10(t1875);let-99(RNAi) | 242.5 | 92.9 | 8 | vs ced-10(t1875) | 0.0044 |
| | | | | vs let-99(RNAi) | 0.0913 |

Table S5. Statistics corresponding to data presented in Fig. 5

| **Fig. 5A Furrowing onset (seconds after NEB)** | | | | | |
|---|---|---|---|---|---|
| **Condition** | **Mean** | **SD** | **n** | **ANOVA, Šídák *** | |
| | | | | **comparison** | **p-value** |
| WT | 178.2 | 10.8 | 11 | N/A | N/A |
| let-99(dd17) | 217.5 | 18.9 | 8 | vs WT | 0.0002 |
| ani-1(RNAi) | 181.5 | 17.2 | 13 | vs WT | 0.9864 |
| let-99(dd17);ani-1(RNAi) | 259.2 | 23.9 | 12 | vs WT | <0.0001 |
| | | | | vs let-99(dd17) | <0.0001 |
| **Fig. 5B Centration at NEB (% Embryo Length)** | | | | | |
| **Condition** | **Mean** | **SD** | **n** | **ANOVA, Šídák *** | |
| | | | | **comparison** | **p-value** |
| WT | 48.2 | 2.5 | 11 | N/A | N/A |
| let-99(dd17) | 59.3 | 3.6 | 10 | vs WT | <0.0001 |
| ani-1(RNAi) | 49.4 | 2.5 | 13 | vs WT | 0.8138 |
| let-99(dd17);ani-1(RNAi) | 59.6 | 3.3 | 12 | vs WT | <0.0001 |
| | | | | vs let-99(dd17) | 0.9988 |
| **Fig. 5C Spindle angle at NEB (degrees)** | | | | | |
| **Condition** | **Mean** | **SD** | **n** | **ANOVA, Šídák *** | |
| | | | | **comparison** | **p-value** |
| WT | 11.8 | 14.4 | 11 | N/A | N/A |
| let-99(dd17) | 40.0 | 25.7 | 10 | vs WT | 0.0021 |
| ani-1(RNAi) | 20.8 | 13.0 | 13 | vs WT | 0.6022 |
| let-99(dd17);ani-1(RNAi) | 30.5 | 16.8 | 12 | vs WT | 0.0500 |
| | | | | vs let-99(dd17) | 0.6164 |
| **Fig. 5D Posterior centrosome position at NEB + 160s (% Embryo Length)** | | | | | |
| **Condition** | **Mean** | **SD** | **n** | **ANOVA, Šídák *** | |
| | | | | **comparison** | **p-value** |
| WT | 79.1 | 2.0 | 11 | N/A | N/A |
| let-99(dd17) | 72.9 | 2.4 | 10 | vs WT | <0.0001 |
| ani-1(RNAi) | 80.7 | 1.6 | 13 | vs WT | 0.2588 |
| let-99(dd17);ani-1(RNAi) | 73.9 | 2.6 | 12 | vs WT | <0.0001 |
| | | | | vs let-99(dd17) | 0.7624 |
| **Fig. 5E Spindle length at NEB + 160s (microns)** | | | | | |
| **Condition** | **Mean** | **SD** | **n** | **ANOVA, Šídák *** | |
| | | | | **comparison** | **p-value** |
| WT | 21.8 | 1.9 | 11 | N/A | N/A |
| let-99(dd17) | 17.2 | 1.1 | 10 | vs WT | <0.0001 |
| ani-1(RNAi) | 22.0 | 1.9 | 13 | vs WT | 0.9962 |
| let-99(dd17);ani-1(RNAi) | 17.3 | 1.2 | 12 | vs WT | <0.0001 |
| | | | | vs let-99(dd17) | 0.9994 |
| **Fig. 5G Extrusions per embryo (number)** | | | | | |
| **Condition** | **Mean** | **SD** | **n** | **ANOVA, Šídák *** | |
| | | | | **comparison** | **p-value** |
| WT | 0 | 0 | 12 | N/A | N/A |
| let-99(dd17) | 0.5 | 1.1 | 8 | vs WT | 0.4476 |
| ani-1(RNAi) | 0.5 | 0.7 | 11 | vs WT | 0.4534 |
| let-99(dd17);ani-1(RNAi) | 1.8 | 0.9 | 12 | vs ani-1(RNAi) | 0.0002 |
| | | | | vs let-99(dd17) | 0.0010 |
| **Fig. 5H Persistence of extrusions (seconds)** | | | | | |
| **Condition** | **Mean** | **SD** | **n** | **ANOVA, Šídák *** | |
| | | | | **comparison** | **p-value** |
| let-99(dd17) | 90.0 | 14.1 | 4 | N/A | N/A |
| ani-1(RNAi) | 50.0 | 17.3 | 5 | N/A | N/A |
| let-99(dd17);ani-1(RNAi) | 90.9 | 57.1 | 24 | vs ani-1(RNAi) | 0.2092 |
| | | | | vs let-99(dd17) | 0.9993 |

* Ordinary one-way ANOVA followed by Šídák's multiple comparisons test

**Supplemental Movies –**

All movies are oriented with anterior to the left. Original movies were taken at either 1 frame per 5 secs or 1 frame per 10 sec as indicated, but playback speed has been adjusted so that all play at same speed; timestamp is relative to NEB=0.

**Movie 1_N2_control** (10-second frame interval).
Movie corresponds to embryo used for the control in Figs 1,3-5.

**Movie 2_let-99(RNAi)** (5-second frame interval).
Movie corresponds to embryo used for Fig. 4; embryo has time to furrowing onset in the control range, but exhibits persistent protrusions near the furrow.

**Movie 3_ced-10(t1875) (**10-second frame interval).
Movie corresponds to embryo used for Fig. 4; embryo exhibits excess cortical activity of the AB cell; small bleb structures are also visible early in the one-cell stage and again later at the two-cell.

**Movie 4_ ced-10(t1875) let-99(RNAi) (**10-second frame interval).
Movie corresponds to embryo used for Fig. 4; embryo has time to furrowing onset in the control range, but exhibits persistent protrusions near the furrow after the first division. Small bleb structures are also visible before NEB of the one-cell stage.

**Movie 5_let-99(dd17) (**10-second frame interval).
Movie corresponds to embryo used for Fig. 5; embryo has delayed furrow onset and exhibits polar protrusions at the anterior membrane and one persistent protrusion near the furrow.

**Movie 6 _ani-1(RNAi) (**10-second frame interval). Movie corresponds to embryo used for Fig. 5; embryo exhibits a posterior polar extrusion during anaphase and protrusions near the furrow after division.

**Movie 7_ ani-1(RNAi); let-99(dd17)** (10-second frame interval).
Movie corresponds to embryo used for Fig. 5; embryo exhibits an anterior polar extrusion during anaphase that travels from the bottom around the pole before resolving, and a second anterior extrusion that travels from the top around the pole. Embryo also exhibits a persistent furrow protrusion.
